# Color compensation in anomalous trichromats assessed with fMRI

**DOI:** 10.1101/2020.08.18.256701

**Authors:** Katherine E.M. Tregillus, Zoey J. Isherwood, John E. Vanston, Stephen A. Engel, Donald I.A. MacLeod, Ichiro Kuriki, Michael A. Webster

## Abstract

Anomalous trichromacy is a common form of congenital color-deficiency resulting from a genetic alteration in the photopigments of the eye’s light receptors. The changes reduce sensitivity to reddish and greenish hues, yet previous work suggests that these observers may experience the world to be more colorful than their altered receptor sensitivities would predict, potentially indicating an amplification of post-receptoral signals. However, past evidence suggesting such a gain adjustment rests on subjective measures of color appearance or salience. We directly tested for neural amplification by using fMRI to measure cortical responses in color-anomalous and normal control observers. Color contrast response functions were measured in two experiments with different tasks to control for attentional factors. Both experiments showed a predictable reduction in chromatic responses for anomalous trichromats in primary visual cortex. However, in later areas V2v and V3v, chromatic responses in the two groups were indistinguishable. Our results provide direct evidence for neural plasticity that compensates for the deficiency in the initial receptor color signals and suggest that the site of this compensation is in early visual cortex.

## Introduction

Typical color perception in humans is trichromatic, beginning with responses to light from three types of cone photoreceptors, known as long-, medium-, and short-wave receptors (L, M, and S) according to the wavelengths they are most sensitive to. A second stage of processing compares signals from the cones; for example the L and M cone responses are subtracted from each other in post-receptoral cells that receive opposing inputs (excitation or inhibition) from them. However, in common inherited color deficiencies the L or M photopigment gene is absent or altered [1-3], leading to a loss of one receptor type (dichromacy), or to a shift in wavelength sensitivity of the affected cone so that it is more similar to the normal M or L cone (anomalous trichromacy). The latter results in a smaller difference in the L vs M signal, and produces weaker sensitivity to the colors conveyed by this difference [4-7]. As an X-linked recessive trait, L vs M color deficiencies affect 6-8% of people with XY chromosomes, but are relatively rare in those with XX (<1%).

Color deficiencies are often modeled as a reduced form of normal trichromacy, in which visual processing is the same except for an initial alteration in the photopigments. In dichromats this reduction model accounts well for poorer color detection and discrimination thresholds in color deficient observers, and has played a critical role in estimating the cone spectral sensitivities [8, 9]. It also is the basis for most simulations of color deficiencies [10-12]. However, in anomalous trichromats the relationship between cone spectra and chromatic sensitivity can be highly variable and complex [13]. Moreover, for these observers the reduction model does not consider potential plasticity in visual coding, which could in principle compensate for a loss in sensitivity by amplifying the post-receptoral difference signals. In anomalous trichromats, the weakened L vs M signal could be restored by gain changes within color-opponent mechanisms, analogous to turning up the contrast on a monitor [14, 15]. Testing for such amplification provides an ideal natural model for studying the limits of long-term compensation in the visual system.

Yet despite this theoretical importance, whether and how color compensation occurs in anomalous trichromacy remains poorly understood (for review see [13] and [16]. The tasks indicating compensation tend to involve judging the similarity or size or salience of color differences [17-19], and thus could potentially reflect conceptual rather than perceptual adaptations. There have been few direct tests of compensation by measuring neural activity [20], and to our knowledge none using functional MRI to assess color processing in anomalous trichromats, a technique that has been shown to be an effective measure of color and contrast coding throughout cortex [21-26] [27-30]. Multiple color-selective areas have been identified along the ventral pathway [31-33], each with potentially different functions and representations of chromatic information. We focused on characterizing the strength of the responses to chromatic contrast in early cortical areas. Primary visual cortex appears to show some plasticity with respect to color over both the long [34] and short term [35], the latter consistent with numerous psychophysical and physiological studies, including fMRI [36, 37]. Thus, if amplification occurs in the neural responses to color for anomalous observers, then early visual cortex is a likely candidate site.

## Results

### Detection Thresholds

Thresholds were collected for 5 out of the 7 anomalous trichromats (AT), and 6 of the 7 color normal controls (CN) (3 observers were unavailable for the threshold session). As expected, the ATs showed dramatically higher thresholds for detecting stimuli defined by L vs M cone differences. On average L vs M thresholds were 7.3 times higher for AT than control participants (AT mean = 9.39, AT SD = 1.81, CN mean = 1.29, CN SD = 0.25, t(9) = 10.98, p<0.001), while thresholds for detecting S vs LM stimuli, which should be relatively unaffected by the color deficiency, did not significantly differ (S vs LM color axis; AT mean = 1.77, AT SD = 0.35, CN mean = 1.71, CN SD = 0.57, t(9) = 0.221, P = 0.830)

### Experiment 1: Simple Fixation Task

We conducted two fMRI experiments to assess neural responses to chromatic contrast in the AT and CN observers. In the first, data were collected from a total of 10 participants (5 CN, 2 deuteranomalous, and 3 protanomalous AT), and stimuli consisted of reversing radial sinewave gratings along the 2 chromatic axes presented at 4 contrast levels (Figure 2A). Participants monitored the fixation point for occasional changes (from black to white) to ensure they were attending to the display. fMRI responses to the gratings arose most strongly in early visual areas, and the amount of activation was similar across groups, with greater chromatic responses in the ventral portions of V2 and V3 than in other regions, consistent with previous work [38]. Accordingly, we used the ventral portions of V2 and V3 in all analyses (Figure 2B, 3).

**Figure 1.**
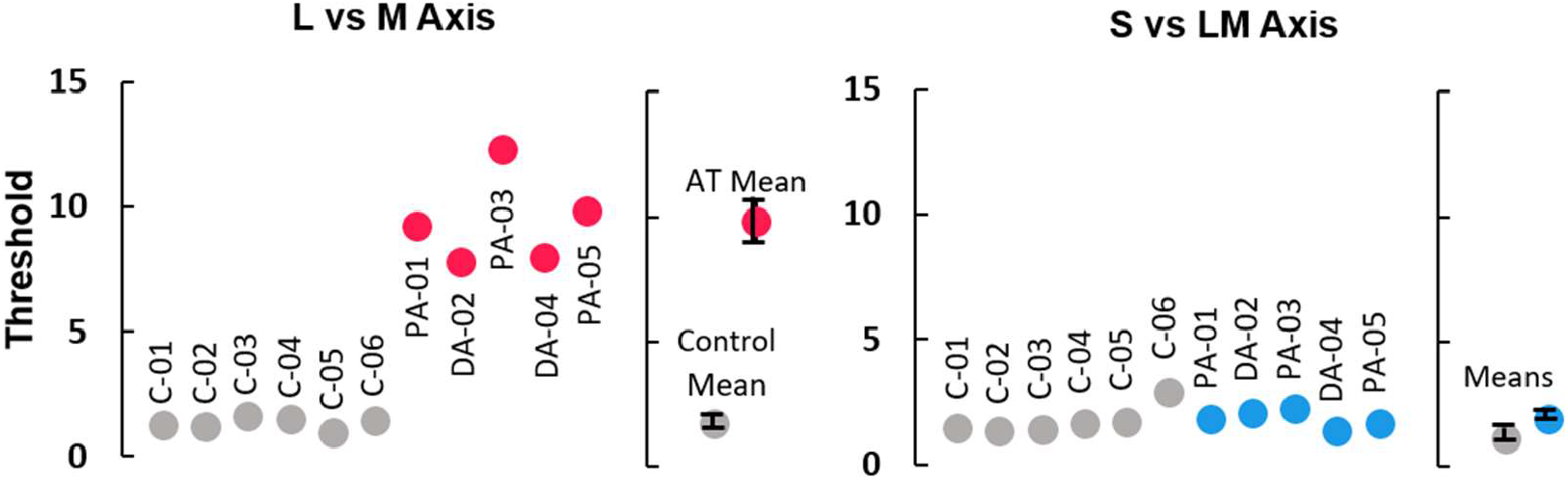
Detection Thresholds. Chromatic contrast detection thresholds for the two chromatic axes, represented in nominal multiples of threshold where thresholds for observers with no color deficits should be at about 1. Settings are shown for each individual observer and for the mean of the control (gray) or anomalous (red or blue) participants. Error bars represent one standard error of the mean. (See also Table S3 for individual participant summaries.)

**Figure 2.**
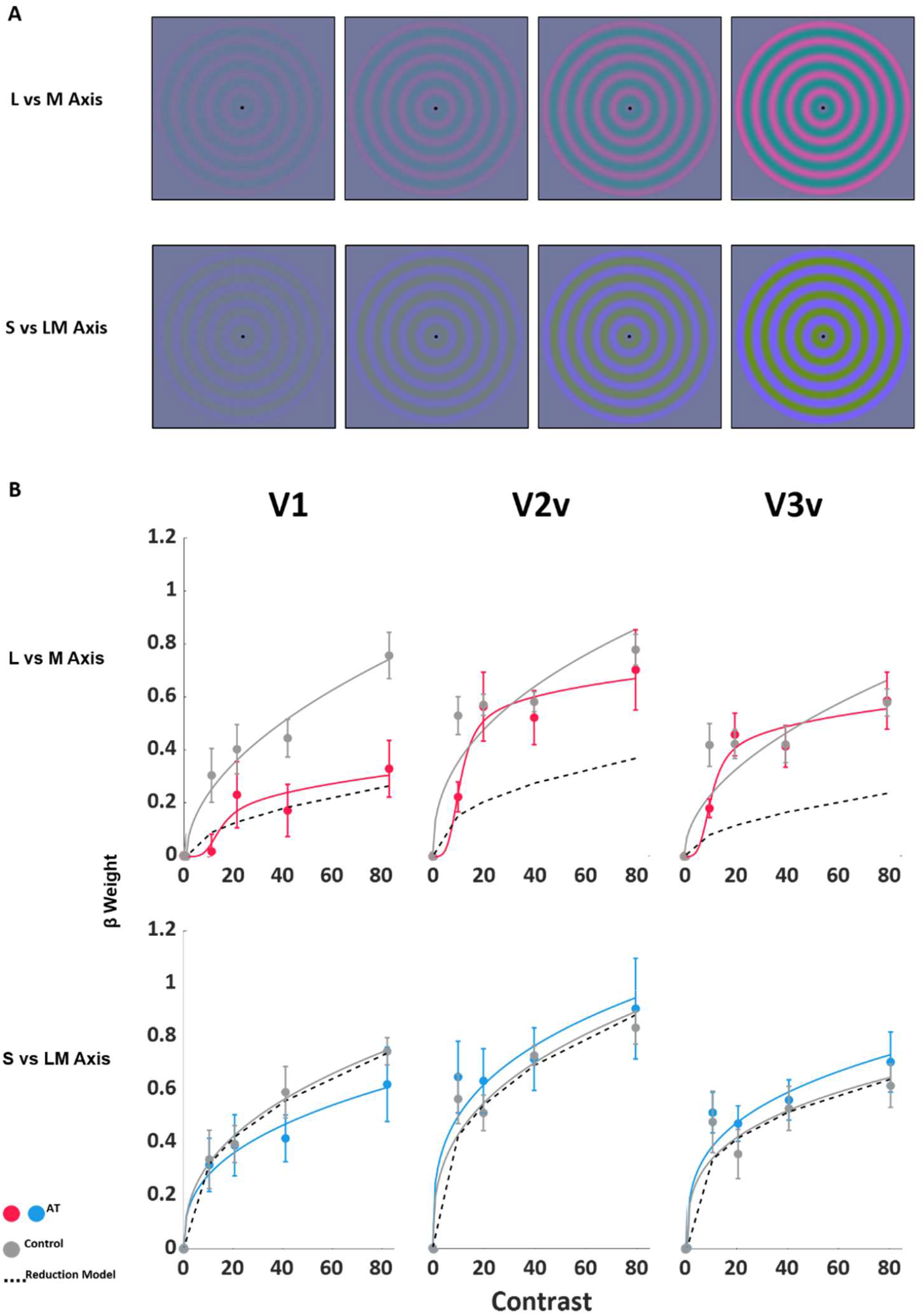
Experiment 1: Simple Fixation Task. Results from Experiment 1. A) Illustration of the 2 color axes and 4 contrast levels used for both experiments. B) Points are β weights from Exp. 1, representing the estimated amplitude of the fMRI response during the simple fixation task. The top row are responses to the L vs M stimuli, and the bottom row are responses to the S vs LM stimuli. Columns are the three regions-of-interest: V1 (left), V2v (center), and V3v (right). The solid red and blue lines are model contrast response functions (CRFs) fitted to the data. The black dashed lines are response predictions for the AT group using the reduction model (see section on Testing for Amplification). Error bars are standard error of the mean. (See also Table S1, Figure S2, and Figure S3 for full statistics and individual results.)

The fMRI data showed reduced responses to L vs M stimuli for AT observers compared to normals in V1, but strikingly this reduction disappeared in V2 and V3, consistent with amplification in these later areas (Figure 2B). The statistical reliability of this pattern was assessed with a 2 x (3 × 2 x 4) mixed ANOVA using the β weights calculated from the GLM analysis of the fMRI data. The between subject factor was *Observer Type* (CN, AT), and the three within subject factors were *Visual Area* (V1, V2v, V3v), *Color Axis* (L vs M, S vs LM), and *Contrast* (10, 20, 40, 80 in nominal threshold units). Post-hoc pairwise comparisons were performed for the L vs M data in each ROI. All statistics reported were Greenhouse-Geisser corrected for sphericity, and pairwise comparisons were Bonferroni corrected for multiple comparisons. Critically, there was a significant difference between ATs and CNs (CI 95%, p = 0.04) in the L vs M response in V1, but not in V2v (CI 95%, p = 0.62) or V3v (CI 95%, p = 1.00). In comparison AT and CN responses were relatively equal in all three areas for S vs LM contrast, where amplification was not predicted. This overall pattern was evidenced in the ANOVA as a highly significant interaction between Group, Visual Area, and Color Axis (F_*3*.*195,25*.*558*_ = 3.67, *p* = 0.023,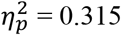). See Table S1 for full ANOVA results.

### Experiment 2: High-Attentional Fixation

A second experiment was designed to replicate our results using a more demanding attentional task. Attention and task can strongly modulate fMRI responses and alter the relationship between response amplitude and stimulus contrast [39, 40], as well as the representation of color, especially in later visual areas [28]. In this experiment we used a highly-demanding fixation task to further isolate “bottom-up” processing to the unattended color patterns. As in past work, observers monitored a stream of black and white numbers at fixation for specific feature conjunctions (e.g. a white 3) [41]. Measurements were collected for 5 AT and 7 CN observers, 3 AT participants and 5 controls participated in both Experiment 1 and 2.

The fMRI data revealed a very similar pattern to that observed in Experiment 1, again supporting amplification – with weaker L vs M responses in V1 for AT observers, but not in later regions (Figure 3, V1: CI 95%, p = 0.02, V2v: CI 95%, p = 0.995, V3v: CI 95%, p = 1.00), see Table S1 for full ANOVA results. The closeness of the two curves suggests that amplification almost completely compensated for receptoral differences by bringing AT cortical responses to CN levels. As in Experiment 1, there were no reliable differences between the ATs and CNs in cortical responses to S vs LM contrasts. As a further control, data were also collected from 1 protanopic observer (male, 31). This participant did not show consistent contrast response functions to the L vs M stimuli, thus providing support that our stimuli were specifically targeting chromatic channels (see Figure S4).

**Figure 3.**
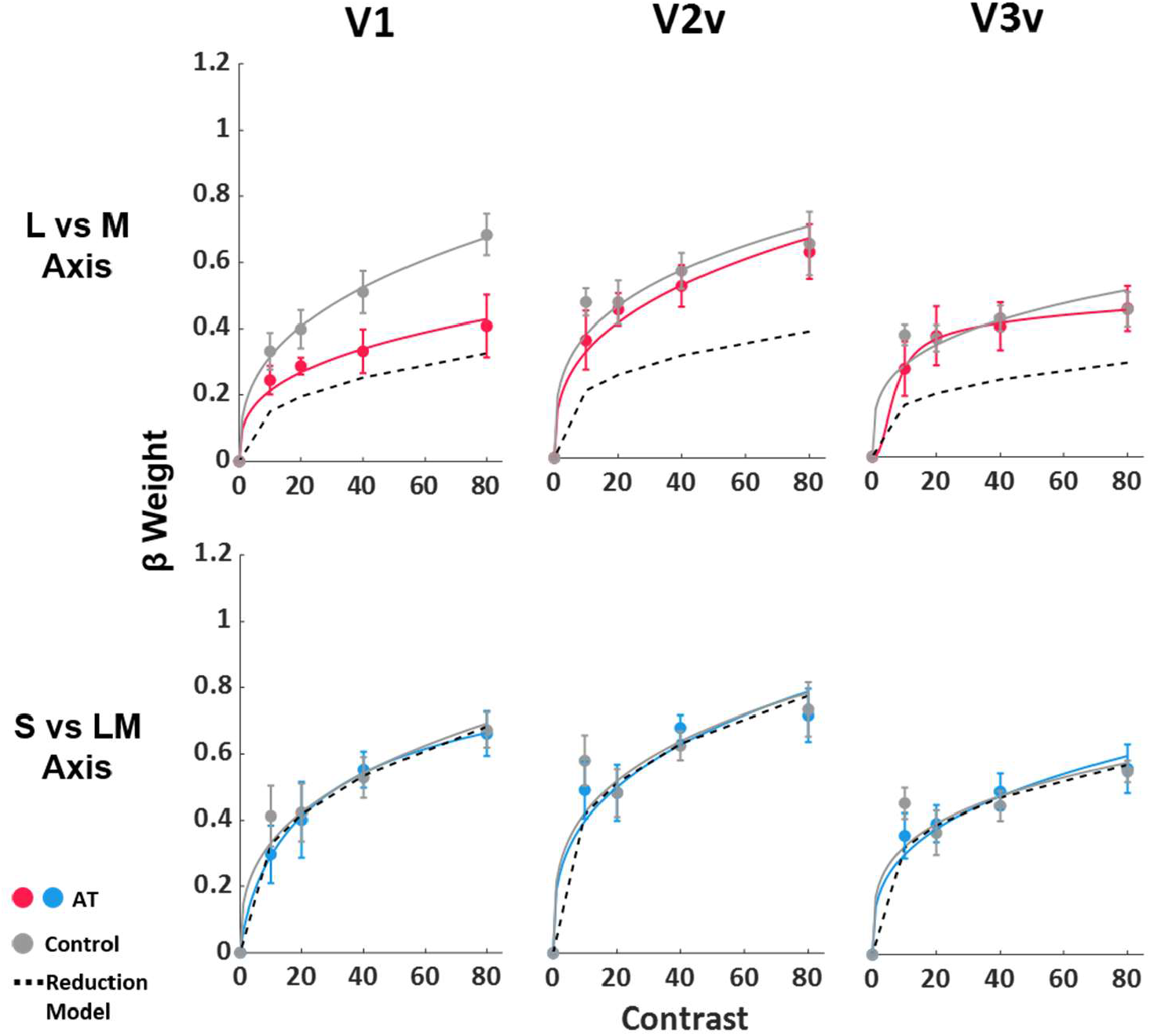
Experiment 2: High-Attention Fixation Task. Results from Experiment 2, high-attention fixation task for the L vs M axis. Plotting conventions are as in Figure 2. (See also Tables S1 and S2 and Figures S1, S2, and S3 for full statistics, individual results, and comparisons between the two conditions.)

5 CNs and 3 ATs participated in both Experiments 1 and 2. We used their data to compare the two attention tasks. (See Figure S1). Overall, AT participants showed slightly lower activation in general during the simple fixation task, but this difference was not significant and only appeared in area V1. We also found no significant effects of the attention task (F_*1,6*.*000*_ = 3.74, *p* = 0.101, 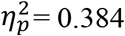). (See Table S2 for full ANOVA.)

### Testing for Amplification by Modeling Contrast Response

To quantify post-receptoral gain in the chromatic signal, we compared cortical responses to an implementation of the reduction model. As a first step, we quantified how responses varied with contrast, by fitting to the data a standard contrast response function (CRF) that has been previously used to quantify the changes in BOLD signal with contrast [42]:

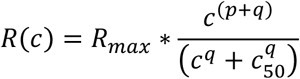

where R is the measured BOLD response, c is the stimulus contrast, and Rmax, c50, p, and q control the function’s shape. Rmax, c50, p, and q were all allowed to vary. The fitted functions are shown as the solid lines in Figures 2B and 3, with control observers in gray, and AT observers in red and blue, and these fitted functions accounted for the data well. Note that these fits were calculated separately for each visual area, since the contrast response functions tend to flatten in higher visual areas [43-46]. See Table S4 for fitted parameters.

We next compared responses of AT observers to predictions of the reduction model, which assumes that all differences between groups are due to photoreceptor sensitivity differences (i.e. that no amplification occurs). The model assumes: 1) that L vs M sensitivity differences arise when information from the cones is first combined into color-opponent signals, that include L vs M contrast, 2) that this reduced L vs M signal propagates through the entire visual pathway, and 3) that the reduced L vs M signal strength is proportional to the reduction in L vs M contrast detection thresholds. This model predicts that L vs M differences at threshold index an identical loss in effective contrast in each area along the cortical pathways, and so CRFs for AT observers in each region should be the same as CNs after rescaling the effective stimulus contrast by the ratio of normal to anomalous thresholds. This rescaling produces a shift of the curve along the horizontal contrast axis, and is a standard model of reduced visual sensitivity (e.g. it is also routinely used to account for differences in sensitivity to different spatial frequencies, velocities, etc.) The model predictions are shown as the black dashed line in Figures 2B and 3., and they approximate the weaker BOLD responses observed in V1 but clearly fall below the responses in V2v and V3v. (See Figures S2 and S3 for individual data and fits.), suggestive of amplification in the AT observers.

To quantify the amount of amplification, we instead scaled the stimulus contrast (c) to fit each AT observer’s individual CRF, with other parameters held constant, and compared this to the contrast scaling predicted by the reduction model (i.e. scaled by their thresholds). Note again that this comparison was done separately for each area to control for differences in the CRF across areas. A ratio of 1 would indicate that the AT’s CRF was consistent with the reduction model, while values greater than 1 would indicate compensation. The resulting ratios again showed clear evidence for amplification. Mean values were 2.94 (sd = 2.81) for V1, 6.39 (sd = 5.21) for V2v, and 7.82 (sd = 5.76) for V3v. For Experiment 2, where we had threshold measures for all 5 AT observers, we compared this scaling factor to 1 (the reduction model prediction), and it was significantly above 1 in V2v (t(4) = 5.12, p = 0.01) and in V3v (t(4) = 4.27, p = 0.03), but not in V1 (t(4) = 1.42, p = 0.07). We also performed a repeated measures ANOVA on the scaling factors and found a significant effect of ROI, (F(2) = 4.98, p = 0.04).

## Discussion

Our results point to strong compensation for color losses in anomalous trichromacy through amplification of cortical responses to chromatic contrast. This in turn suggests that the greater than expected responses to colors observed behaviorally [17-19] may reflect actual gain changes in early cortical processing. Both the neural and behavioral evidence are not inconsistent with the profound sensitivity losses in color-anomalous observers in threshold discrimination for color, since these thresholds are limited by noise, which may also be amplified by the gain change [14, 15, 47]. The suprathreshold compensation for color contrast may be similar to effects observed for spatial contrast, where thresholds depend strongly on spatial frequency (pattern size) yet once above threshold perceived contrast is independent of frequency [48]. In this regard it is notable that less compensation is evident at the lowest contrast we tested (10) which was near the mean threshold for the anomalous observers, though this difference only appears in Experiment 1.

Our results also point to an early but not initial stage of cortical coding as the site of the contrast amplification. Both experiments showed that the L vs M responses were consistent with the reduction model in V1 but were indistinguishable from color-normals in V2v and V3v. The fact that this pattern persisted under the attentionally demanding task of Experiment 2 suggests top-down modulation is not responsible for the neural gain occurring between V1 and V2v. Given the plasticity of V1, it is not clear why a long-term adjustment for chromatic contrast would first emerge in V2. While chromatic processing is known to undergo a series of transformations along the ventral stream, the function of different areas and representations remains uncertain [27, 30, 49, 50]. How these stages might be impacted by a color deficiency could further elucidate not only the nature of the plasticity but also their potential role in color perception. Thus it would be instructive in future studies to explore the magnitude and pattern of compensations at higher stages of the ventral pathway.

Importantly, however, our results do not preclude a gain change occurring in V1, since the BOLD contrast response could be disproportionately driven by the input layers and thus depend on the responses inherited from the LGN [51]. Thus it may be that AT and CN participants have more similar responses in the output layers of V1 (for review see [52]). Moreover, the differences between V1 and V2v varied for individual observers, consistent with previous studies showing that anomalous trichromats are a highly heterogeneous group [18, 53]. Of our 7 AT participants, 3 seemed to show more similar levels of activation to CN observers in area V1 (Figure S3, Table S3). Considering this, we cannot rule out some amplification within V1. One AT (PA-01) also showed unusually low responses to every level of L vs M contrast (Figure S3). However, the general pattern we observed persisted when the results were re-analyzed without including this observer.

The weak chromatic responses in V1 do imply that there is relatively little compensation prior to cortex. This conclusion is also consistent with a recent study using visual evoked potentials, which found evidence for color compensation but only when the stimuli were viewed binocularly [20], implying a cortical site since signals from the two eyes first converge in V1. The emergence of compensation in the cortex would mean that the retina and LGN remain in a weakened, unadjusted state for processing the L vs M signals. While there are increasing signs of short-term plasticity, specifically contrast adaptation, at these early stages [54-56], contrast adaptation is known to be substantially stronger in the visual cortex [36], especially within the parvocellular pathway along which the L vs M signals are carried [54, 57].

The adjustments we measured could potentially represent a very long-term form of visual plasticity, though we cannot rule out a process that acts more rapidly. Color vision adjusts to changes in the environment or observer over widely varying timescales [58], and compensation for the color losses could similarly involve multiple timescales and processes. For example a recent study found that color percepts in anomalous observers could be enhanced after just a few days wearing glasses that increased the L vs M contrasts [59]. One potential mechanism that could account for the compensatory adjustments we observed is sensory adaptation, in which neural signals adjust to the average and range of the ambient stimulation [60]. If the contrast is too low, then a neuron may increase its sensitivity so that the range of outputs is maintained [47, 61]. Most studies of adaptation have focused on very short-term aftereffects and on sensitivity losses rather than gains. However, exposures to contrast losses as brief as a few hours have been found to lead to compensatory enhancements of contrast sensitivity [62-64]. Our study is a natural color analog of adapting to a contrast reduction but measured over a lifetime of exposure.

The results of the present study are relevant not only to color but to understanding adaptation and neural calibration more generally, for the patterns and mechanisms of plasticity appear to be very similar across sensory levels and modalities [37]. Our results suggest that the nervous system can, at least in some cases, strongly compensate for a weakened initial coding of stimuli. How and under what circumstances such compensation can occur has important implications for treatment of visual disorders, as well as for the increasing prospects of gene therapy for color deficiencies [65]. In turn, it is clear that perception and performance for many visual tasks remains compromised for most anomalous trichromats, and even when there is clear evidence for compensation it is typically not complete [21]. Understanding what factors act to mitigate full compensation might help elucidate fundamental neural constraints on sensory plasticity [16].

## Author Contributions

Conceptualization, K.E.M.T., M.A.W., S.A.E., and D.I.A.W; Methodology, K.E.M.T., M.A.W., S.A.E, D.I.A.M, and I.K.; Data Collection, K.E.M.T. and J.E.V.; Data Analysis, Z.J.I. and K.E.M.T.; Writing – Original Draft, K.E.M.T. and Z.J.I.; Writing – Review & Editing, M.A.W., S.A.E., I.K., D.I.A.M, and J.E.V.

### Acknowledgements

Research supported by EY-10834 and P20-GM103650 (MW). Additional support for color screening and participant recruitment from Dr. Michael Crognale and the Crognale Color Vision Assessment Clinic at UNR.

## Declaration of Interests

The authors declare no competing interests.

## STAR Methods

### Resource Availability

#### Lead Contact

Further information and requests for resources should be sent to the Lead Contact, Katherine E.M. Tregillus. There is no restriction for distribution of materials.

#### Material Availability

This study did not generate any new reagents.

#### Data and Code Availability

Original/source data for Figures 2B and 3 in the paper are available for download on the Open Science Framework at https://osf.io/2sv9y

### Experimental Model and Subject Details

A total of 7 anomalous trichromats (AT) were recruited from The University of Nevada-Reno and the community (3 deuteranomalous, 4 protanomalous, all male), ages 19-42. Color-normal control participants were all UNR students (including author KT), ages 24-38, 2 female. We also tested one dichromat (a protanope) as a further control (see Figure S4). 3 AT participants and 5 controls participated in both Experiment 1 and 2. AT participants were recruited with flyers and classroom announcements specifically seeking “colorblind” participants, or who had failed color screenings from previous assessments at UNR and who had agreed to be contacted for future experiments. Due to our recruitment methods (including initial self-diagnosis), it is likely we gathered participants with generally more severe deficits. Chromatic contrast detection thresholds were collected for 6 of the controls and 5 AT participants, 2 AT and 1 CN were not tested because they were not available. All procedures were approved by the University of Nevada, Reno’s institutional review board, and all participants provided informed consent prior to testing.

### Method Details

#### Anomaloscope

The color vision of all participants was assessed using standard color screening techniques, including Rayleigh matching using a Heidelbert-Multi-Color Anomaloskop (OCULUS, Inc., Wetzlar, Germany). The match range was then used to calculate the anomaly quotient (AQ), which was used as the diagnostic criterion with an AQ of less than 0.7 indicating protanomalous, an AQ range of 0.7 to 1.4 color normal, and an AQ of greater than 1.4 deuteranomalous, where 1 is an equal mixture of red and green, less than 1 is a higher proportion of red, and greater than 1 is a higher proportion of green. Dichromats are able to match the orange test field to almost any red-green mixture by adjusting the brightness, and are thus easily differentiable from AT observers.

#### Stimuli

Stimuli were presented on calibrated computer screens in testing rooms at UNR or the fMRI facility at Renown Health Hospital (Reno, NV). Thresholds were collected using a SONY 20SE monitor, with displays controlled by a Cambridge Research Systems (Kent, UK) Visual Stimulus Generator (VSG) board, which allowed for high color resolution. fMRI stimuli were displayed on a 32 in. SensaVue (85 Hz refresh rate) monitor (Invivo, Inc. Gainesville, FL) situated behind the scanner bore and viewed through a head-mounted mirror. The monitor’s maximum visual field was 31 deg. by 19 deg. Both monitors were calibrated with a Photo Research (Syracuse, NY) PR655 spectroradiometer, and gun outputs were linearized through lookup tables.

Participants were presented with phase reversing (1 Hz) radial sinewave gratings (0.28 c/deg., 14.5 deg field) defined by chromatic variations along either the L vs M or S vs LM cone-opponent axes (Figure 2A). The same stimuli were used in the scanner and during the threshold task, similar to stimuli used by Mullen et al. [26]. Both chromatic directions were shown at 4 levels of chromatic contrast for a total of 8 conditions (Figure 2A). Contrasts were scaled roughly as multiples of detection threshold for normal trichromats based on a previous study [66, 67]. In units of pooled cone contrast the highest contrast stimuli were 12% and 70% for L vs M and S vs LM gratings respectively. All stimuli were produced with the Psychophysics Toolbox [68] for MATLAB (Mathworks Inc., Natick, MA).

#### Isoluminance Settings

Participants were first adapted to a neutral gray background for 1 minute, and then completed a minimum motion task to determine isoluminance [69]. The minimum motion stimuli consisted of alternating achromatic and chromatic concentric sinusoidal gratings that were offset in phase. Participants were instructed to adjust the gratings until the stimuli no longer appeared to radiate inward or outward. Stimulus luminance was adjusted individually for each observer. Individual isoluminance measures were obtained separately for the different display systems used for the threshold and fMRI experiments.

#### Chromatic Detection Thresholds

We used a temporal 2-alternative forced choice task to measure contrast detection thresholds. The grating was shown in one of two temporal intervals (1 sec. on, 1 sec. off) signaled by beeps, and participants pressed a button to indicate at which interval the grating appeared. Contrast was varied in two randomly interleaved staircases (2 up, 1 down). Each staircase terminated after 10 reversals. The average of the last 8 reversals across each staircase was taken as the participant’s detection threshold. S vs LM and L vs M contrast thresholds were collected during the same session, but across different runs. Thresholds were collected in a separate session and location from the fMRI data.

#### fMRI Procedure

Data were acquired on a Philips 3T Ingenia scanner using a 32-channel digital SENSE head coil (Philips Medical Systems, Best, Netherlands). Functional data for Experiments 1 and 2 were obtained using T2 *-weighted, echo planar images (2 sec. TR, 180 volumes, voxel size 2.75 × 2.75 × 3 mm^3,^ 36 slices, 55 ms inter slice time, 0 mm gaps). We used a block design where a run consisted of 16 blocks of 14 sec. each, interleaved with 8 sec. fixation-only gaps. Each stimulus type was presented twice per run, and the order was counterbalanced across runs. Each run lasted 360 sec. In Experiment 1, all participants completed 8 runs. In Experiment 2, participants completed 6 runs, reduced from Experiment 1. Anatomical scans occurred halfway through a session in order to reduce motion artifacts. Anatomical scans were collected using T1 weighted images (voxel size of 1×1×1 mm^3,^ 30 sec transverse slices, 17 ms TE, 76° flip angle, and 220 × 220 mm^2^ field of view). Anatomical and functional runs were collected within a single session for 12 of the participants, while the remaining 2 participants completed functional runs across two sessions.

#### Fixation Tasks

During scans in Experiment 1, participants completed a simple fixation task to ensure that they were responsive and fixating during the stimulus presentation. The black fixation circle flickered at random, jittered intervals, and the participants were instructed to make a response when a change occurred. In order to further test and control for attentional effects on the BOLD contrast response, Experiment 2 used a more demanding fixation task [39]. The fixation mark for this task was a number that randomly switched from black to white at a jittered rate, while also changing to a random number from 1 to 9. Participants were assigned a random pair of number conjunctions and asked to press a button when either conjunction appeared (e.g. black nine or white three) [41].

### Quantification and Statistical Analysis

#### fMRI Data Analysis

Images acquired from the scanner were pre-processed before analysis. This included slice scan time correction as well as motion correction, which were both conducted using SPM12 (SPM software package, Wellcome Department, London, UK; http://www.fil.ion.ucl.ac.uk/spm/) and custom lab scripts. Data were motion corrected using a rigid body transform and 7th degree B-spline interpolation. Images were slice scan time corrected (ascending, interleaved) using the first image as the reference slice and resliced into the space of the first image.

Volumetric segmentation of white matter was performed using Freesurfer (v6.0) (http://surfer.nmr.mgh.harvard.edu/) [70, 71]. 3D surface reconstructions of the left and right hemisphere were generated using mrMesh (a function within the mrVista Toolbox) by growing a 3-voxel thick layer above the gray/white boundary. To improve data visualization (i.e. when projecting functional data onto surfaces), these surfaces were also computationally-inflated using the “smoothMesh” option in mrMesh (8 iterations). Note that the cortical surface models were only used for data visualization and region-of-interest (ROI) definition. All analyses and statistics were performed using the volumetric data.

Due to limited scan time, an anatomical template of retinotopic maps in early visual areas was used to approximate the cortical location of visual areas (V1, V2, V3) and eccentricity ranges that the stimulus subtended (0.00°–9.50°) [72, 73]. The retinotopy template was fitted to each subject’s anatomical segmentation using Freesurfer (v6.0) (http://surfer.nmr.mgh.harvard.edu/) [70, 71]. This is a well validated approach to estimating the location of early visual areas that accounts for anatomical variability across subjects by aligning and morphing the template onto each individual subject’s anatomy (see Figure 4A for an example of the retinotopic template aligned to a subject’s cortical surface). As a sanity check, we plotted percent signal change from ROIs drawn from the template (see Figure 4B for an example), and we find that ROIs across visual areas V1, V2v, and V3v demonstrate reliable signal change in response to each color/contrast condition used in the present study.

**Figure 4.**
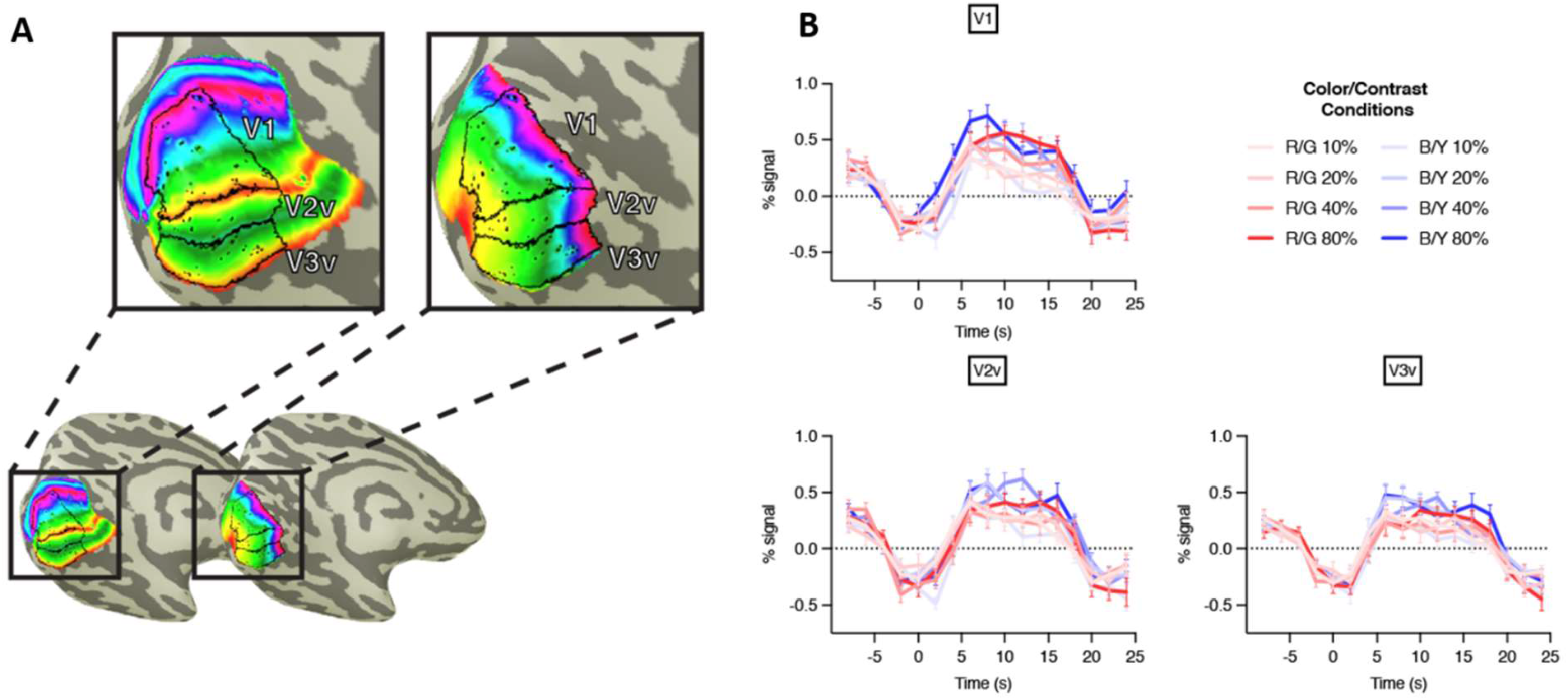
fMRI Data Analysis. A) Example retinotopic template (polar angle–left, eccentricity—right) aligned to an exemplar subject’s left hemisphere (sub-04, CN, exp-02). V1, V2v, and V3v ROIs are drawn on the inflated cortical surface depicted here. The range of the eccentricity map was clipped to 0.00° to 9.50°, corresponding to the range the stimulus subtended. The ROIs have the central 0.00°–0.95° clipped to exclude the fixation point and the mean luminance gray gap between the fixation point and the stimulus. B) Plots of percent signal change in V1, V2v, and V3v in an exemplar subject (sub-04, CN, exp-02). The data was averaged across 6 runs. Each run consisted of 16 blocks where each condition was presented twice (6 runs x 2 repetitions = 12 repetitions total for each condition). The stimulus was presented for 14 seconds (stimulus onset is at 0 seconds in the plots above). A mean luminance blank screen was presented for 8 seconds between blocks.

#### Statistics

Analysis of the fMRI results were performed using SPSS (IMB Corp., Chicago, IL). A 2 x (3 × 2 x 4) mixed ANOVA was conducted on the β weights calculated from the GLM analysis of the fMRI data for Experiment 1 (n = 10, 5 CN, 5 AT) and Experiment 2 (n = 12, 7 CN, 5 AT). The between subject factor was *Color Vision Type* (CN, AT), and the three within subject factors were *Visual Area* (V1, V2v, V3v), *Color* (L vs M, S vs LM), and *Contrast* (10, 20, 40, 80). See Results section and Table S1 for these results. To compare Experiment 1 and Experiment 2 results, another ANOVA was conducted with the additional within subject factor of *Attention* (high, low). Only participants in both experiments were included in this analysis (n = 8, 5 CN, 3 AT). See Results section and Table S2 for these results.

#### Fitting Contrast Response Function

Further analysis in the mrVista Toolbox included a general linear model (GLM) of responses across early visual areas (V1, V2v, V3v) for each individual subject. The eccentricity range 0.00°–0.95° was excluded from each ROI prior to analysis to exclude the fixation point and mean luminance gray gap between the fixation point and the stimulus. A standard gamma-function HRF was used to model the hemodynamic response function [42]. All runs were concatenated and the null gray background condition was used as baseline. After β weights were calculated, a repeated measures ANOVA was performed on these values in order to determine the effects of each condition individually across visual areas.

We then fit data with a contrast response function:

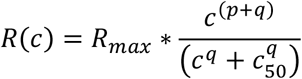

where Rmax, c50, p, and q varied freely, though to aid fitting we set an upper-bound of 4 for p and q since previous work estimates their values for early visual areas to sit between 0.1 and 2.0 [42]. No bounds were set on Rmax and c50. In its typical form, when c>c50, the function is a simple power function with an exponent of p+q. The function is compressive at low contrasts, and expansive at high contrasts. See Table S4 for a list of all fitted parameters.

To compute predictions of the reduction model, we scaled the contrast by the ratio of thresholds of CN and AT observers, t:

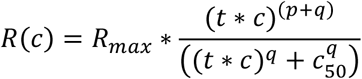

Other values were fixed at values that best fit the mean response from CN observers.

To compute the amount of amplification, we further scaled contrast by a scaling factor, sc, and again kept all other parameters fixed at values from the mean CN response:

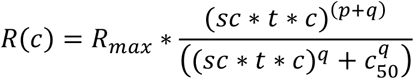

We estimated sc for each individual AT observer, and these results are listed in Table S3. Because it is a multiplicative factor, we used the log of the scaling parameter in a one-sample t-test to determine if the AT responses were amplified, i.e. they differed significantly from their threshold-scaled predictions, which will be the case if the scaling factor differs from 1.

## Supplemental Information

**Figure S1.**
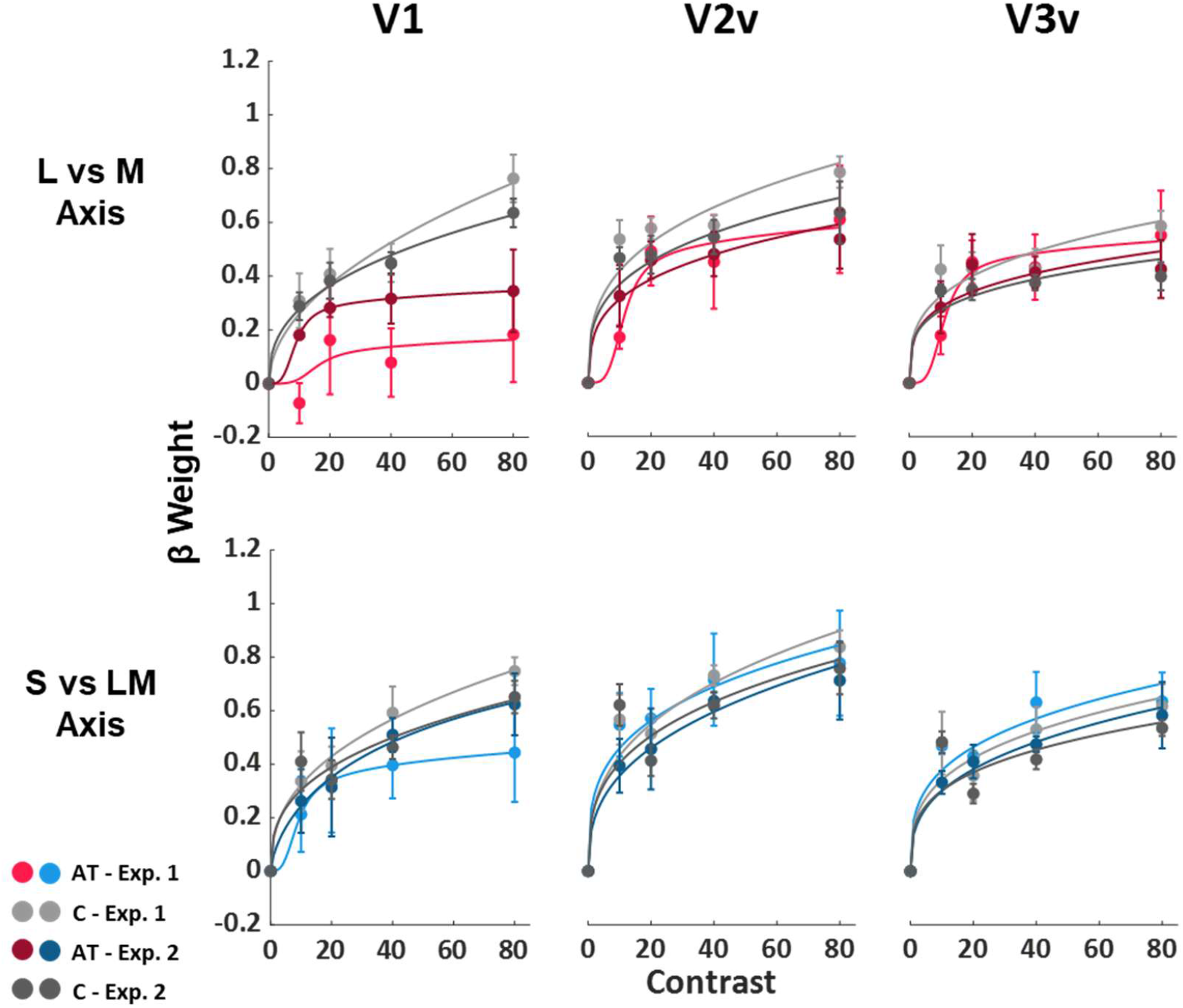
Comparison of Experiment 1 (simple fixation task) and Experiment 2 (high-attention fixation task). Figure shows only data from those who participated in both experiments. ATs are shown in red and blue and CNs are shown in gray. Experiment 1 data is shown in lighter colors. Solid lines are fitted CRFs. **Related to Figures 2B and 3 in the main text and section Method Details in STAR Methods**.

**Figure S2.**
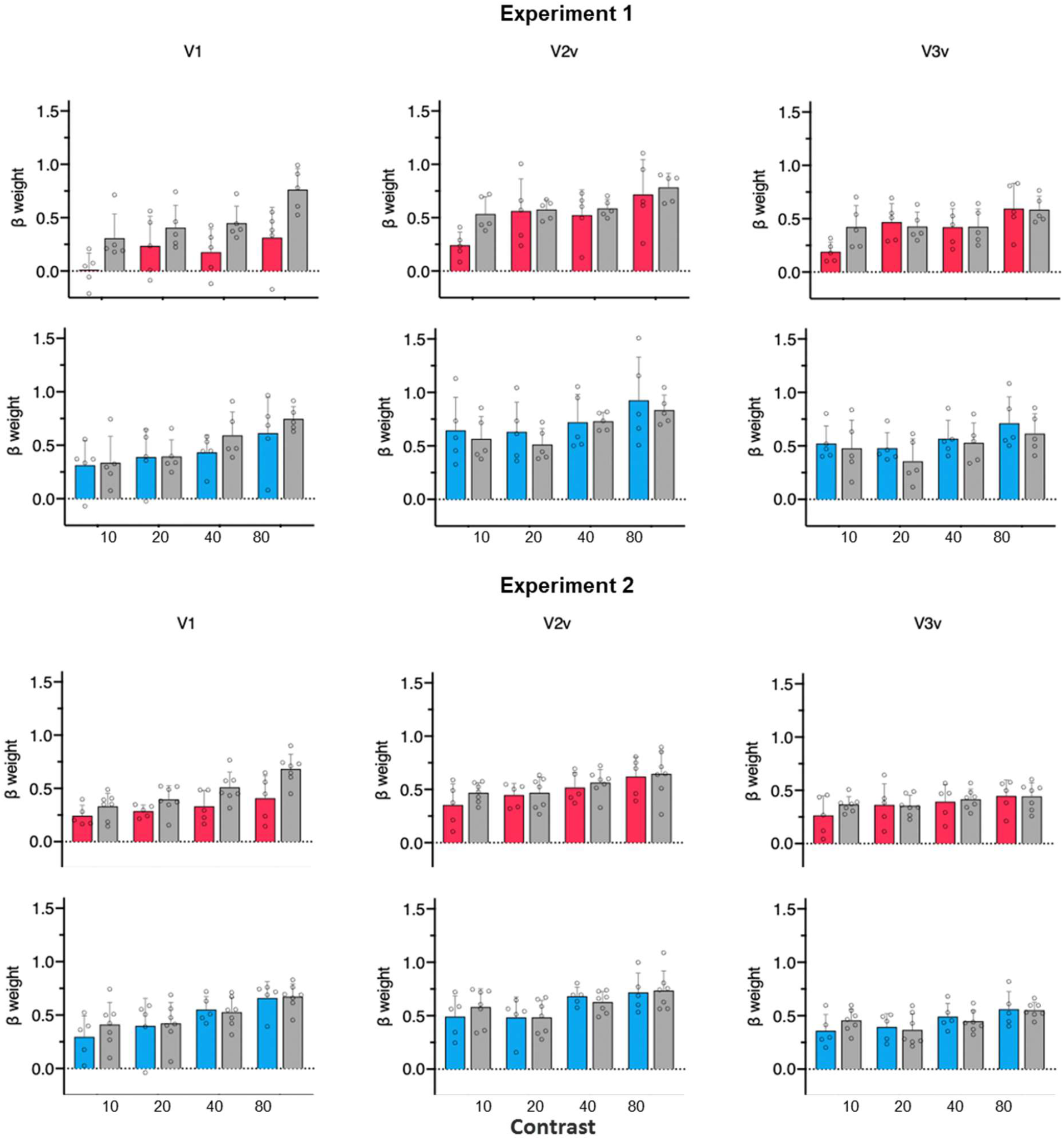
Box plots showing mean data with individual participants’ data shown as open circles. L vs M data are shown in red, and S vs LM data are shown in blue. Error bars represent 1 standard deviation. **Related to Figures 2B and 3 in main text**.

**Figure S3.**
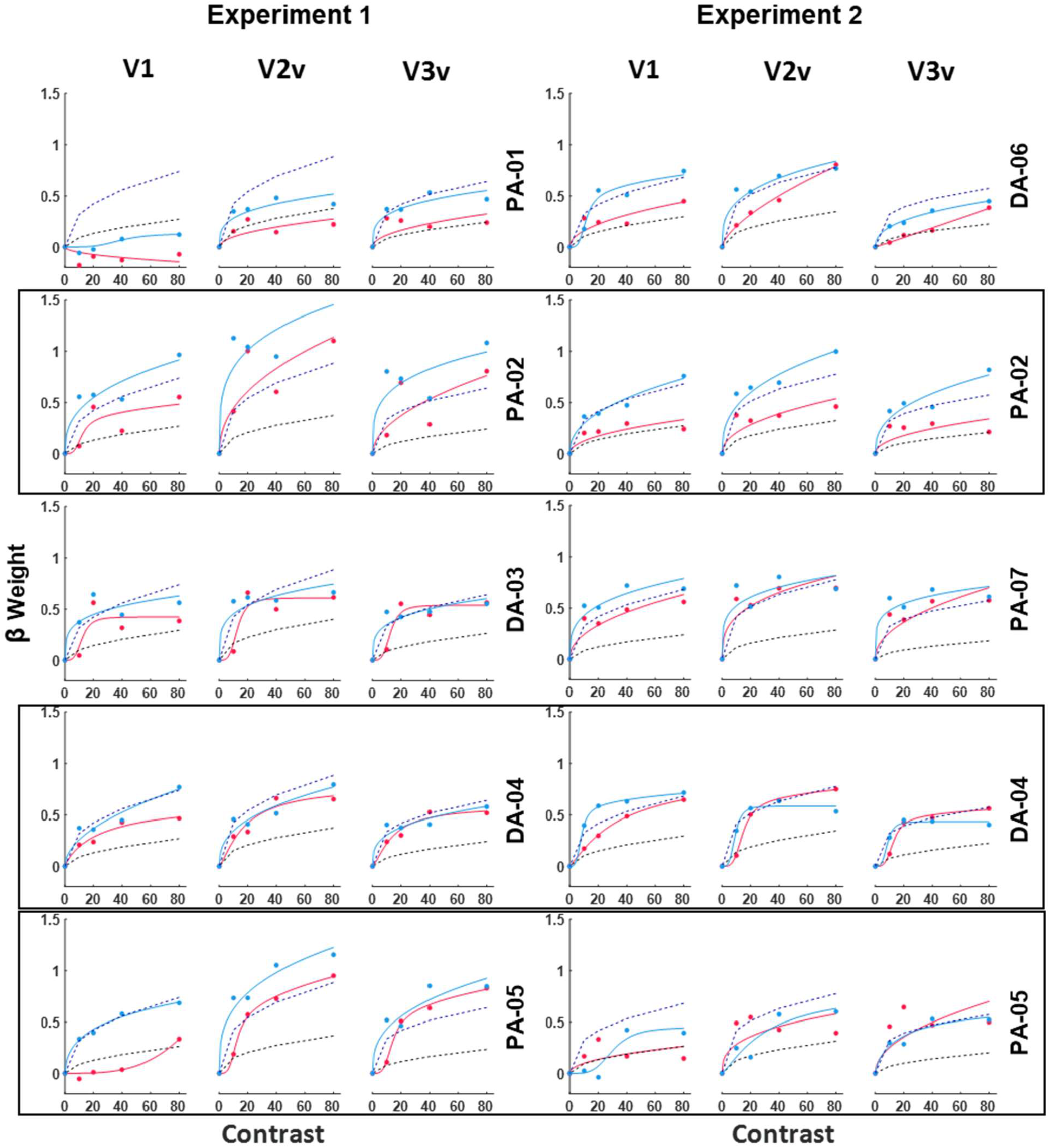
Individual Results. L vs M responses (red) and S vs LM responses (light blue) for each AT observer, for Experiment 1 (simple fixation task) and Experiment 2 (high-attention fixation task). Black dashed lines show the L vs M threshold prediction. Dark blue dashed lines show the S vs LM threshold prediction (a scaling of ∼1 from mean CN responses for all AT participants). Where individual thresholds were unavailable, the mean AT threshold was used. Participants who were in both experiments are outlined in black. **Related to Figures 2B and 3 in main text**.

**Figure S4.**
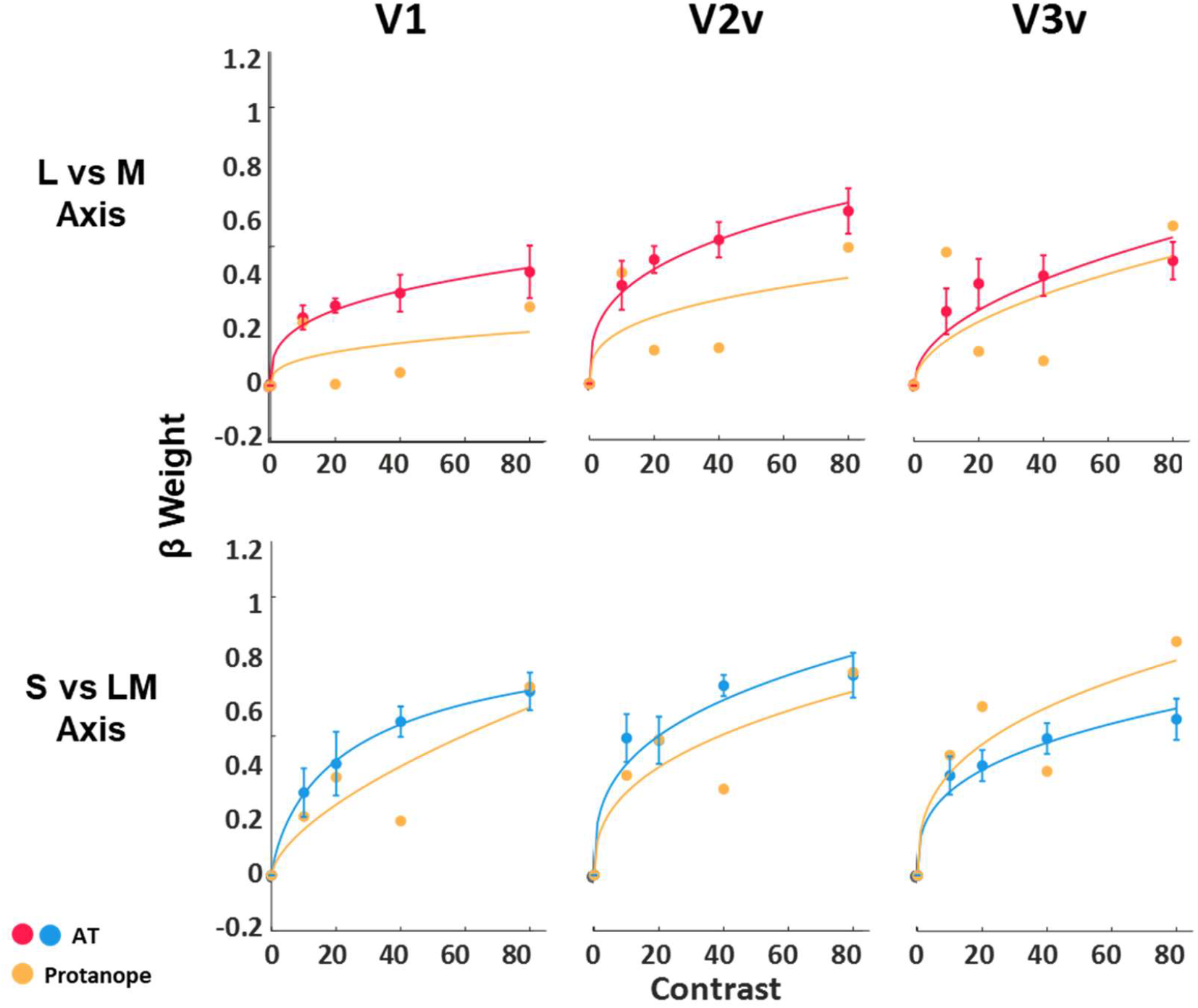
Dichromatic Observer. One dichromatic (protanopic) participant was tested using the fMRI paradigm of Experiment 2 (male, 31). PN-01 was diagnosed as a protanope based on anomaloscope settings, in which he accepted all red-green ratios as matches. BOLD responses to chromatic contrast levels for PN-01 (shown in yellow), compared to the mean activation across AT observers for areas V1, V2v, V3v. L vs M responses are in the top row, and S vs LM responses are in the bottom row. This observer’s thresholds were 9.14 for the L vs. M stimuli and 2.02 for the S stimuli, similar to the thresholds we obtained for the anomalous observers. This observer did not show a monotonic contrast response to the L vs M stimuli, thus providing support that stimuli were specifically targeting chromatic channels. **Related to Figure 3 in main text and section Experimental Model and Subject Details in STAR Methods**.

**Table S1.** Statistics: Experiment 1 and Experiment 2 (V1, V2v, V3v Mixed ANOVA). Summary of ANOVA main effects and interactions for Experiments 1 and 2. **Related to section Quantification and Statistical Analysis in STAR Methods**.

| Experiment 1 |  |  |  |  |  |  |
| --- | --- | --- | --- | --- | --- | --- |
| | df | Error | F | p | $\eta_p^2$ | Power |
| **Visual Area** | **1.710** | **13.677** | **10.578** | **0.002** | **0.569** | **0.947** |
| Visual Area x Color Vision Type | 1.710 | 13.677 | 2.262 | 0.146 | 0.220 | 0.357 |
| **Color** | **1.000** | **8.000** | **21.163** | **0.002** | **0.726** | **0.979** |
| **Color x Color Vision Type** | **1.000** | **8.000** | **10.566** | **0.012** | **0.569** | **0.812** |
| **Contrast** | **2.545** | **20.357** | **19.837** | **0.000** | **0.713** | **1.000** |
| Contrast x Color Vision Type | 2.545 | 20.357 | 1.005 | 0.400 | 0.112 | 0.219 |
| Visual Area x Color | 1.294 | 10.355 | 2.934 | 0.111 | 0.268 | 0.379 |
| Visual Area x Color x Color Vision Type | 1.294 | 10.355 | 2.195 | 0.167 | 0.215 | 0.296 |
| **Visual Area x Contrast** | **3.195** | **25.558** | **3.531** | **0.027** | **0.306** | **0.733** |
| **Visual Area x Contrast x Color Vision Type** | **3.195** | **25.558** | **3.672** | **0.023** | **0.315** | **0.752** |
| **Color x Contrast** | **2.500** | **19.997** | **6.401** | **0.005** | **0.444** | **0.898** |
| Color x Contrast x Color Vision Type | 2.500 | 19.997 | 2.960 | 0.065 | 0.270 | 0.565 |
| **Visual Area x Color x Contrast** | **2.905** | **23.240** | **4.332** | **0.015** | **0.351** | **0.795** |
| **Visual Area x Color x Contrast x Color Vision Type** | **2.905** | **23.240** | **4.092** | **0.019** | **0.338** | **0.769** |
| Experiment 2 |  |  |  |  |  |  |
| **Visual Area** | **1.844** | **18.443** | **8.563** | **0.003** | **0.461** | **0.923** |
| Visual Area x Color Vision Type | 1.844 | 18.443 | 0.731 | 0.484 | 0.068 | 0.151 |
| **Color** | **1.000** | **10.000** | **8.246** | **0.017** | **0.452** | **0.735** |
| Color x Color Vision Type | 1.000 | 10.000 | 1.211 | 0.297 | 0.108 | 0.169 |
| **Contrast** | **1.980** | **19.804** | **19.905** | **<0.001** | **0.666** | **1.000** |
| Contrast x Color Vision Type | 1.980 | 19.804 | 0.764 | 0.478 | 0.071 | 0.161 |
| Visual Area x Color | 1.449 | 14.491 | 0.879 | 0.403 | 0.081 | 0.157 |
| **Visual Area x Color x Color Vision Type** | **1.449** | **14.491** | **5.909** | **0.020** | **0.371** | **0.715** |
| **Visual Area x Contrast** | **3.170** | **31.704** | **8.177** | **<0.001** | **0.450** | **0.987** |
| Visual Area x Contrast x Color Vision Type | 3.170 | 31.704 | 2.084 | 0.119 | 0.172 | 0.496 |
| Color x Contrast | 2.361 | 23.611 | 0.510 | 0.637 | 0.049 | 0.129 |
| Color x Contrast x Color Vision Type | 2.361 | 23.611 | 0.370 | 0.729 | 0.036 | 0.106 |
| Visual Area x Color x Contrast | 2.589 | 25.893 | 1.967 | 0.150 | 0.164 | 0.417 |
| **Visual Area x Color x Contrast x Color Vision Type** | **2.589** | **25.893** | **3.635** | **0.031** | **0.267** | **0.690** |
| *Notes: Reported statistics were adjusted using Greenhouse-Geisser correction. Rows in bold represent interactions with $p < 0.05$.* |  |  |  |  |  |  |

**Table S2.**
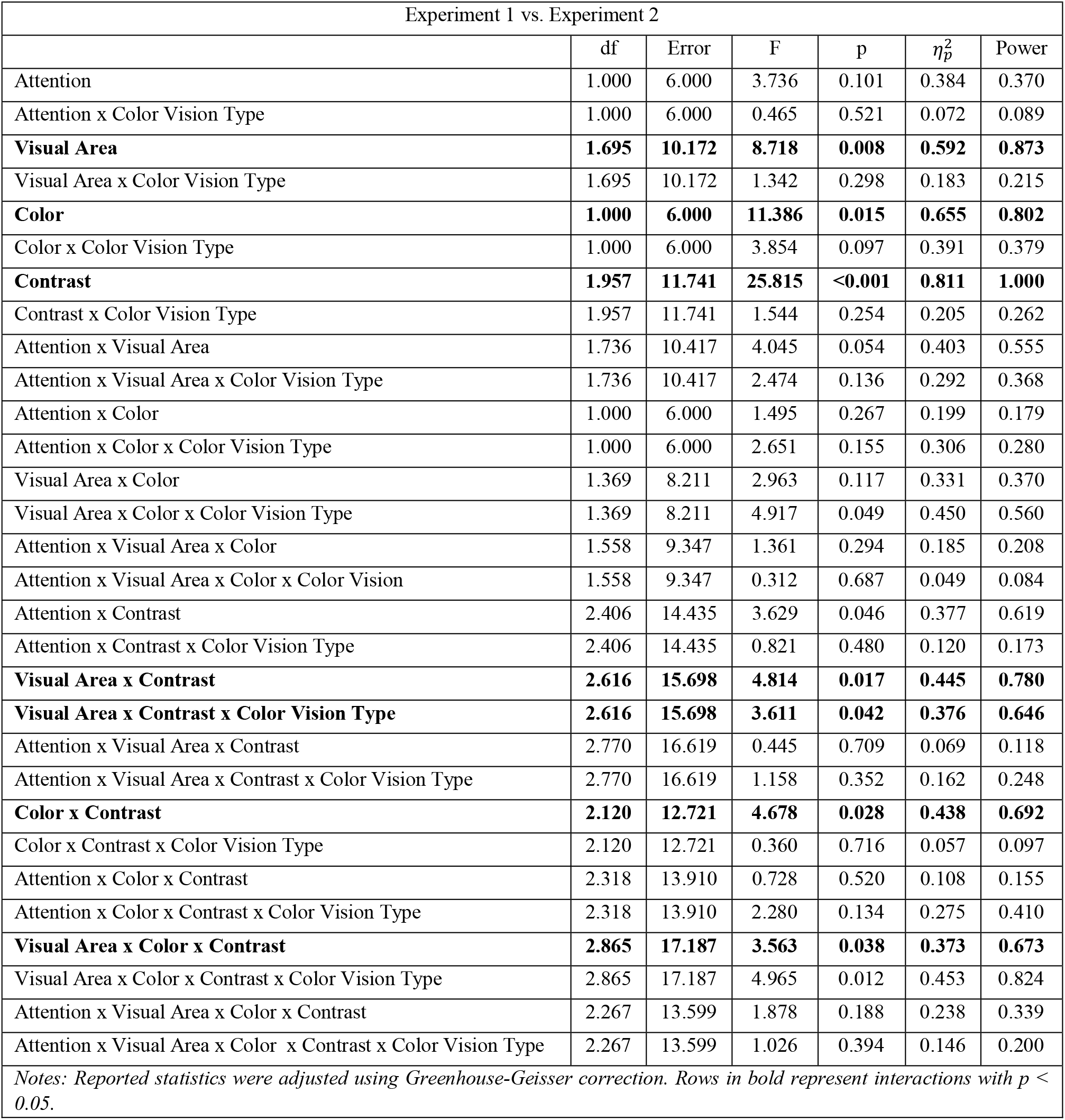
Statistics: Experiment 1 vs Experiment 2 (V1, V2v, V3v Mixed ANOVA). Summary of ANOVA main effects and interactions for comparison between the attentional effects of Experiment 1 and Experiment 2. **Related to section Quantification and Statistical Analysis in STAR Methods**.

| Experiment 1 vs. Experiment 2 |  |  |  |  |  |  |
| --- | --- | --- | --- | --- | --- | --- |
| | df | Error | F | p | $\eta_p^2$ | Power |
| Attention | 1.000 | 6.000 | 3.736 | 0.101 | 0.384 | 0.370 |
| Attention x Color Vision Type | 1.000 | 6.000 | 0.465 | 0.521 | 0.072 | 0.089 |
| **Visual Area** | **1.695** | **10.172** | **8.718** | **0.008** | **0.592** | **0.873** |
| Visual Area x Color Vision Type | 1.695 | 10.172 | 1.342 | 0.298 | 0.183 | 0.215 |
| **Color** | **1.000** | **6.000** | **11.386** | **0.015** | **0.655** | **0.802** |
| Color x Color Vision Type | 1.000 | 6.000 | 3.854 | 0.097 | 0.391 | 0.379 |
| **Contrast** | **1.957** | **11.741** | **25.815** | **<0.001** | **0.811** | **1.000** |
| Contrast x Color Vision Type | 1.957 | 11.741 | 1.544 | 0.254 | 0.205 | 0.262 |
| Attention x Visual Area | 1.736 | 10.417 | 4.045 | 0.054 | 0.403 | 0.555 |
| Attention x Visual Area x Color Vision Type | 1.736 | 10.417 | 2.474 | 0.136 | 0.292 | 0.368 |
| Attention x Color | 1.000 | 6.000 | 1.495 | 0.267 | 0.199 | 0.179 |
| Attention x Color x Color Vision Type | 1.000 | 6.000 | 2.651 | 0.155 | 0.306 | 0.280 |
| Visual Area x Color | 1.369 | 8.211 | 2.963 | 0.117 | 0.331 | 0.370 |
| Visual Area x Color x Color Vision Type | 1.369 | 8.211 | 4.917 | 0.049 | 0.450 | 0.560 |
| Attention x Visual Area x Color | 1.558 | 9.347 | 1.361 | 0.294 | 0.185 | 0.208 |
| Attention x Visual Area x Color x Color Vision | 1.558 | 9.347 | 0.312 | 0.687 | 0.049 | 0.084 |
| Attention x Contrast | 2.406 | 14.435 | 3.629 | 0.046 | 0.377 | 0.619 |
| Attention x Contrast x Color Vision Type | 2.406 | 14.435 | 0.821 | 0.480 | 0.120 | 0.173 |
| **Visual Area x Contrast** | **2.616** | **15.698** | **4.814** | **0.017** | **0.445** | **0.780** |
| **Visual Area x Contrast x Color Vision Type** | **2.616** | **15.698** | **3.611** | **0.042** | **0.376** | **0.646** |
| Attention x Visual Area x Contrast | 2.770 | 16.619 | 0.445 | 0.709 | 0.069 | 0.118 |
| Attention x Visual Area x Contrast x Color Vision Type | 2.770 | 16.619 | 1.158 | 0.352 | 0.162 | 0.248 |
| **Color x Contrast** | **2.120** | **12.721** | **4.678** | **0.028** | **0.438** | **0.692** |
| Color x Contrast x Color Vision Type | 2.120 | 12.721 | 0.360 | 0.716 | 0.057 | 0.097 |
| Attention x Color x Contrast | 2.318 | 13.910 | 0.728 | 0.520 | 0.108 | 0.155 |
| Attention x Color x Contrast x Color Vision Type | 2.318 | 13.910 | 2.280 | 0.134 | 0.275 | 0.410 |
| **Visual Area x Color x Contrast** | **2.865** | **17.187** | **3.563** | **0.038** | **0.373** | **0.673** |
| Visual Area x Color x Contrast x Color Vision Type | 2.865 | 17.187 | 4.965 | 0.012 | 0.453 | 0.824 |
| Attention x Visual Area x Color x Contrast | 2.267 | 13.599 | 1.878 | 0.188 | 0.238 | 0.339 |
| Attention x Visual Area x Color x Contrast x Color Vision Type | 2.267 | 13.599 | 1.026 | 0.394 | 0.146 | 0.200 |
| Notes: Reported statistics were adjusted using Greenhouse-Geisser correction. Rows in bold represent interactions with $p < 0.05$. | | | | | | |

**Table S3.** **Summary of individual values for each AT participant. Related to sections Experimental Model and Subject Details, Method Details, and Quantification and Statistical Analysis in the STAR Methods**.

| Participant | Exp. | CVD Type | L-M Threshold | AQ Match Range | Iso-luminance Adjustment* | CRF Scaling Factor |  |  |  |  |  |
| --- | --- | --- | --- | --- | --- | --- | --- | --- | --- | --- | --- |
|  |  |  |  |  |  | Experiment 1 |  |  | Experiment 2 |  |  |
|  |  |  |  |  |  | V1 | V2v | V3v | V1 | V2v | V3v |
| PA-01 | 1 | Protanomalous | - | 17.6 | 0.085 | - | - | - | - | - | - |
| PA-02 | 1 & 2 | Protanomalous | 9.19 | 5.5 | 0.020 | 3.59 | 8.74 | 9.61 | 1.48 | 2.65 | 2.71 |
| DA-03 | 1 | Deutanomalous | - | 20.8 | -0.033 | - | - | - | - | - | - |
| DA-04 | 1 & 2 | Deutanomalous | 7.92 | 0.04 | -0.032 | 3.55 | 3.60 | 6.09 | 4.65 | 5.01 | 7.48 |
| PA-05 | 1 & 2 | Protanomalous | 9.82 | 5.1 | -0.048 | 0.53 | 6.36 | 12.35 | 0.96 | 3.51 | 12.09 |
| DA-06 | 2 | Deutanomalous | 7.80 | 7.1 | -0.050 | - | - | - | 2.20 | 4.13 | 1.83 |
| PA-07 | 2 | Protanomalous | 12.25 | 20.7 | -0.007 | - | - | - | 7.31 | 8.97 | 15.33 |
| * Isoluminance adjustment is in units of degree of tilt outside of the isoluminant plane, with positive numbers suggesting an upward tilt of the +L axis. Mean CN adjustment = -0.005, no participants were adjusted by more than +/-0.1 degrees. Reported values are specific to the MR-safe display used for Experiments 1 and 2. |  |  |  |  |  |  |  |  |  |  |  |

**Table S4.**
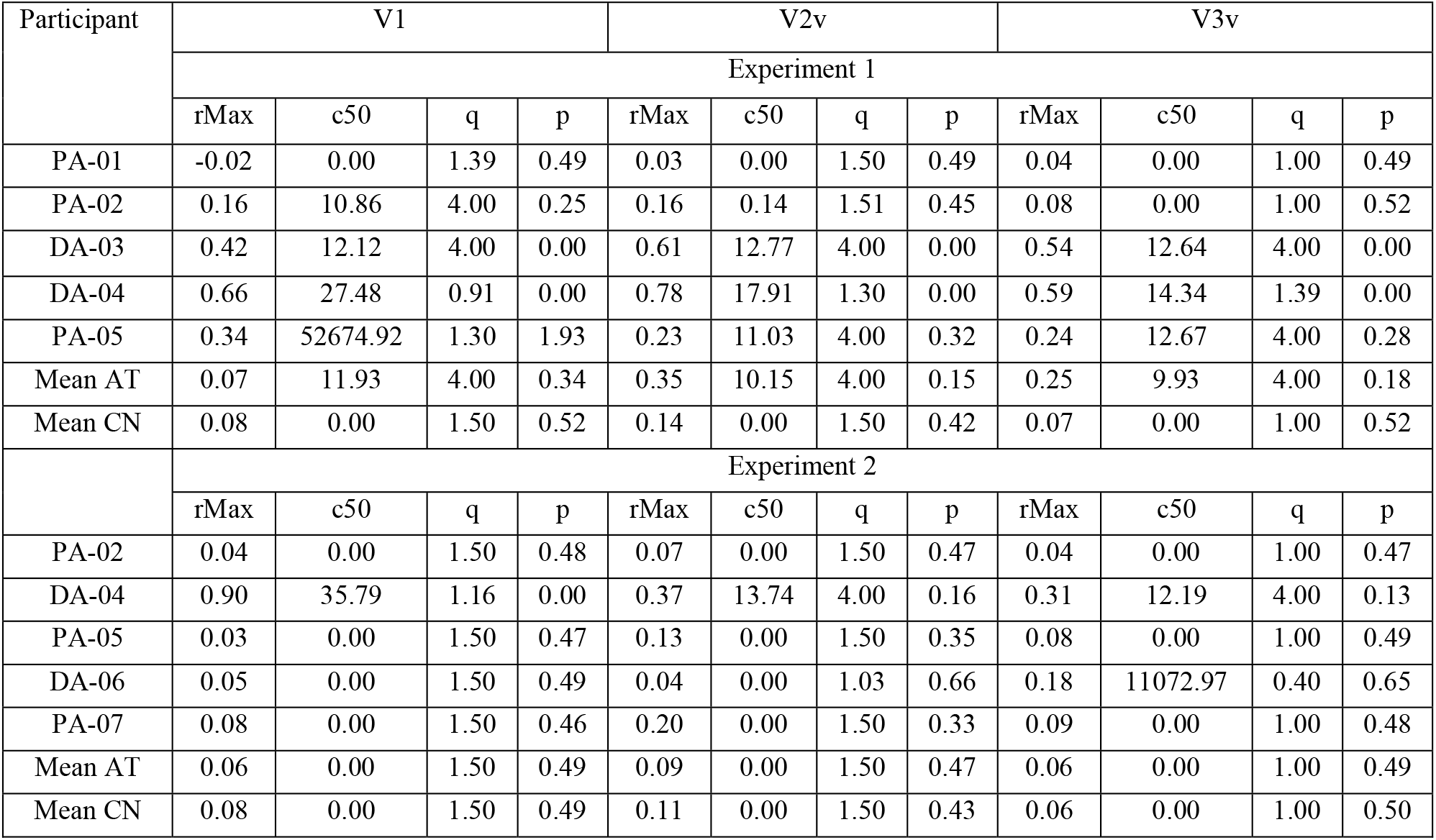
**Summary of individual parameter estimates for each AT and means for AT and CN groups. Related to section Quantification and Statistical Analysis in STAR Methods**.

## Notes

https://osf.io/2sv9y

### Competing Interest Statement

The authors have declared no competing interest.

### Summary of Updates

Revised in response to peer review. Currently in press.

